# Hypoxia-inducible factor (HIF) activation promotes osteogenic transition of valve interstitial cells and accelerates heart valve calcification in chronic kidney disease (CKD)

**DOI:** 10.1101/2023.02.08.527686

**Authors:** Dávid Máté Csiki, Haneen Ababneh, Andrea Tóth, Gréta Lente, Árpád Szöőr, Anna Tóth, Csaba Fillér, Tamás Juhász, Béla Nagy, Enikő Balogh, Viktória Jeney

**Author notes:** The first two authors contributed equally to this study and share first authorship. **Correspondence:** Viktória Jeney, MTA-DE Lendület Vascular Pathophysiology Research Group, Research Centre for Molecular Medicine, Faculty of Medicine, University of Debrecen, Debrecen, Hungary.

## Abstract

**Aims:** Valve calcification (VC) is a widespread complication in chronic kidney disease (CKD) patients on hemodialysis. VC is an active process with the involvement of *in situ* osteogenic transition of valve interstitial cells (VICs). VC is accompanied by the activation of hypoxia inducible factor (HIF) pathway, but the role of HIF activation in the calcification process remains undiscovered.

**Methods and result:** Using *in vitro* and *in vivo* approaches we addressed the role of HIF activation in osteogenic transition of VICs and CKD-associated VC. Elevation of osteogenic (Runx2, Sox9) and HIF activation markers (HIF-1α and HIF-2α) and VC occurred in adenine-induced CKD mice. High phosphate (Pi) induced upregulation of osteogenic (Runx2, alkaline-phosphatase, Sox9, osteocalcin) and hypoxia markers (HIF-1α, HIF-2α, Glut-1), and calcification in VICs. Down-regulation of HIF-1α and HIF-2α inhibited, whereas further activation of HIF pathway by hypoxic exposure (1% O_2_) or hypoxia mimetics (desferrioxamine, CoCl2, Daprodustat (DPD)) promoted Pi-induced calcification of VICs. Pi augmented the formation of reactive oxygen species (ROS) and decreased viability of VICs, whose effects were further exacerbated by hypoxia. N-acetyl cysteine inhibited Pi-induced ROS production, cell death and calcification under both normoxic and hypoxic conditions. DPD treatment corrected anemia but promoted VC in the CKD mice model.

**Conclusions:** HIF activation plays a fundamental role in Pi-induced osteogenic transition of VICs and CKD-induced VC. The cellular mechanism involves stabilization of HIF-1α and HIF-2α, increased ROS production and cell death. Targeting the HIF pathways may thus be investigated as a therapeutic approach to attenuate VC.

**Translational perspective:** One in four hemodialysis-dependent CKD patients on DPD treatment experience a major cardiovascular event during a 2.5-year follow-up period. This work provides a possible explanation for this phenomenon and should initiate further studies to address whether DPD-mediated acceleration of valve calcification triggers the unbeneficial effect of DPD.

## 1. INTRODUCTION

Vascular calcification and valvular heart disease are highly prevalent in patients with chronic kidney disease (CKD). In particular, the prevalence of valve calcification (VC) is eight times higher in end stage renal disease patients undergoing hemodialysis than in the general population^1^. Aortic and mitral valves are affected most frequently, and calcification of both valves arises 10-20 years sooner in CKD patients compared with subjects with normal kidney function^1–3^. Hyperphosphatemia is a critical etiopathogenic factor in CKD-associated vascular and VC^4–6^.

Heart valves are avascular, though metabolically active tissues, composed of an outer monolayer of valve endothelial cells and several internal layers of valve interstitial cells (VICs)^7^. For a long time VC was considered as a passive deposition of calcium-phosphate which supposition was challenged by studies showing the existence of osteoblast-like and osteoclast-like cells in human aortic valve leaflets^8^. About 13% of aortic valves removed during valve replacement surgery contain lamellar bone-like organized structures^9^.

Many lines of evidence suggest that VC is an actively regulated process in which *in situ* phenotypic transition of VICs into osteoblast-like cells and myofibroblasts occurs^7,10–12^. Studies indicated that excessive formation of reactive oxygen species (ROS) play a crucial role in the initiation and progression of these processes^13^. The osteogenic transition of VICs is characterized by elevated expression of osteogenic markers including runt-related transcription factor 2 (Runx2), bone morphogenetic protein 2 (BMP2), alkaline phosphatase (ALP), osteopontin (OPN) and osteocalcin (OCN) ^14,15^. Importantly, these osteogenic markers are found to be upregulated along with increased ROS production in calcified human aortic valves^14–16^.

Adequate nutrition and oxygenation of VICs are ensured through diffusion from the circulation. In case of valve thickening diffusional oxygen transfer is compromised leading to tissue hypoxia. Hypoxic response is mediated by heterodimeric hypoxia inducible transcription factors (HIF)^17^. The HIF pathways are regulated by the oxygen-sensitive alpha subunits which are stabilized and translocated to the nucleus, form heterodimers with the beta subunits and bind to hypoxia response elements in the promoter regions of hypoxia-regulated genes under hypoxic conditions^17^. HIF activation is an adaptive response that helps cell survival and restoration of tissue homeostasis in hypoxic conditions. Systemic response to hypoxia aims to improve oxygen delivery to tissues through dilatation of peripheral vessels and constriction of pulmonary vasculature, and the induction of angiogenesis and erythropoiesis. Whereas cellular response to hypoxia aims to balance ATP production and consumption via increasing the efficiency of anaerobic energy-producing metabolic pathways such as glycolysis, and by decreasing processes with high energy demand such as protein synthesis ^17,18^.

Previous studies showed increased expression of HIF-1α and vascular endothelial growth factor (VEGF) accompanied by neo-angiogenesis in stenotic, thickened and calcified valves, suggesting the activation of the HIF pathway upon valve remodeling^19–24^. Hypoxia and sustained HIF activation have been shown to promote vascular smooth muscle cells (VSMCs) phenotype switch towards osteoblast-like cells, and accelerate vascular calcification^25–27^.

Therefore, in this work we have investigated whether hypoxia and HIF signaling are actively participated in osteogenic trans-differentiation of VICs and subsequent VC. We choose the adenine+high phosphate-induced CKD model as our *in vivo* approach and inorganic phosphate (Pi)-induced calcification of human VICs for the *in vitro* experiments.

## 2. METHODS

### 2.1. Materials

We purchased all the reagents from Sigma-Aldrich Co (St. Louis, MO, USA) unless indicated otherwise.

### 2.2. Induction of CKD and DPD treatment in mice

Mice were kept in plastic cages with standard beddings and unlimited access to food and water. We performed the experiments with the approval of the Institutional Ethics Committee of University of Debrecen under a registration number of 10/2021/DEMÁB, and all procedures conformed to the guidelines from Directive 2010/63/EU of the European Parliament on the protection of animals used for scientific purposes. Animal studies were reported in compliance with the ARRIVE guidelines.

Ten male C57BL/6 mice (8-10 weeks old, n=5/group) were randomly divided into 2 groups: control (Ctrl) and CKD. CKD was induced by an adenine-containing diet as described previously^27,28^. In the first 6 weeks the mice received a diet containing adenine (0.2%) and elevated phosphate (0.7%) followed by adenine (0.2%) and high phosphate (1.8%) diet (S8106-S075 and S8893-S006 respectively, Ssniff, Soest, Germany) for 4 weeks.

In a separate experiment we tested the effect of the hypoxia mimetic drug Daprodustat DPD (HY-17608, MedChemExpress, NJ, USA) on calcification. To this end, 15 male C57BL/6 mice (8-10 weeks old) were divided into 3 groups (Ctrl, CKD, CKD+DPD, n=5/group). DPD was suspended in 1% methylcellulose and was administered orally at a dose of 15 mg/kg/day between weeks 7 and 10 as described previously^27^. The dose of DPD is the minimal dose that corrects anemia in C57BL/6 mice which was chosen based on our previous study^27^. We euthanized the mice by CO_2_ inhalation at the end of the experiments. We collected blood by cardiac puncture for analysis.

### 2.3. Laboratory analysis of renal function and anemia in mice

Plasma phosphate, urea and creatinine levels were assessed spectrophotometrically and by a kinetic assay respectively, on a Cobas^®^ 6000 device (Roche Diagnostics, Mannheim, Germany). Hematology parameters were determined from citrate-anticoagulated whole blood by a Siemens Advia-2120i analyzer (Siemens, Tarrytown, NY, USA) with the use of 800 Mouse C57BL program of Multi Species software.

### 2.4. Imaging and quantification of aortic calcification

OsteoSense™ dye (OsteoSense 680 EX and NEV10020EX; PerkinElmer, MA, USA) was reconstituted in DPBS in a concentration of 20 nmol/mL. We anesthetized the mice with isoflurane inhalation and injected the dye in a dose of 2 nmol/20g body weight retro-orbitally. Imaging was performed 24 hours post-injection. We euthanized the mice with CO_2_ inhalation, perfused with 5 ml of PBS, and analyzed the isolated hearts *ex vivo* by an IVIS Spectrum In Vivo Imaging System (PerkinElmer, MA, USA).

### 2.5. Histology and immunohistochemistry

After the OsteoSense™ imaging, the isolated hearts were fixed in 10% neutral buffered formalin and were embedded in paraffin blocks and cut into 4-5 μm-thick cross-sections. Sections were deparaffinized and rehydrated followed by von Kossa and Alizarin Red stainings with standard procedures. All the sections were counterstained with hematoxylin eosin.

Immunohistochemistry was performed on heart tissue samples to address localization of HIF-1α. Hearts were fixed in 4% paraformaldehyde and embedded in paraffin. Serial sections were made, deparaffinated and rinsed in PBS (pH 7.4). Bovine serum albumin (1%, diluted in PBS) was used to block non-specific binding sites. Sections were incubated with HIF-1α antibody (GTX127309; GeneTex; Irvine, CA, USA) at a dilution of 1:500 at 4°C overnight, followed by anti-rabbit Alexa Fluor 555 secondary antibody (Life Technologies Corporation, Carlsbad, CA, USA) applied at a dilution of 1:1000. As negative control, we applied anti-rabbit Alexa Fluor 555 without the use of primary antibodies. Samples were mounted in Vectashield mounting medium (Vector Laboratories, Peterborough, England) containing DAPI (4’,6-Diamidino-2-phenylindole dihydrochloride) for nuclear DNA staining. Results were obtained by an Olympus FV3000 confocal microscope (Olympus Corporation, Tokyo, Japan) using a 60x PlanApo N oil-immersion objective (NA: 1.42) and FV31S-SW software (Olympus Corporation, Tokyo, Japan). Sequential scan mode was used to recors Z image series of 1-μm optical thickness. For excitation 543 nm laser beam was used. The average pixel time was 4 μsec. Images of Alexa Fluor 555 and DAPI were overlaid using Adobe Photoshop version 10.0 software. Contrast of images was equally increased without changing constant settings.

### 2.6. Cell culture and reagents

Human VICs (P10462, Innoprot, Bizkaia, Spain) were maintained in Fibroblast Medium (P60108, Innoprot) supplemented with 10% FBS (10270-106, Gibco, Grand Island, NY, USA), sodium pyruvate, L-glutamine and antibiotic antimycotic solution, according to the manufacturer’s protocol. Cells were cultured at 37 °C in a humidified atmosphere with 5% CO_2_ content. We performed the experiments on VICs between passages 4 and 8.

To induce calcification we exposed VICs to an osteogenic medium (OM) which was obtained by supplementing the growth medium with inorganic phosphate (Pi in the form of NaH2PO4 and Na2HPO4, pH 7.4, 2.5 mmol/L, or as indicated) and Ca (CaCl2, 0.3 mmol/L).Both growth medium and OM were changed in every other day throughout the experiments.

### 2.7. Hypoxic Treatment

To provide hypoxic environment we placed the cells into a modular incubator chamber (Billups-Rothenberg Inc, Del Mar, CA, USA). We filled the chamber with a gas mixture of 1% O_2_, 5% CO_2_, and 94% of N_2_ (Linde, Dublin, Ireland) and applied a continuous slow flow (0.1 L/min) of the gas throughout the experiment. For normoxia, we used a gas mixture of 21% O_2_, 5% CO_2_, and 74% of N_2_. In other experiments, we used hypoxia mimetic drugs such as desferrioxamine (DFO, 40 μmol/L), CoCl2 (200 μmol/L) and DPD (20 μmol/L) or the HIF-1 inhibitor chetomin (Tocris, Bristol, United Kingdom, 12 nmol/L).

### 2.8. Alizarin Red staining and quantification

At the end of the experiment we washed the cells with PBS, and fixed with 4% paraformaldehyde for 20 minutes. After rinsing with PBS we stained the cells with Alizarin Red S solution (2%, pH 4.2) for 10 minutes at room temperature. Following this we applied several washes with deionized water to remove unbound dye. After taking pictures of the staining, we dissolved the dye in 100 μl of 100 mmol/L hexadecylpyridinium-chloride and determined optical density at 560 nm.

### 2.9. Quantification of Ca deposition

VICs cultured in 96-well plates were washed with PBS and decalcified with HCl for 30 minutes at room temperature. We measured Ca content from HCl-containing supernatants with QuantiChrom Calcium Assay kit (Gentaur, Kampenhout, Belgium). To obtain protein concentration, we washed the cells with PBS and lysed in a lysis buffer containing NaOH (0.1 mol/L) and sodium dodecyl sulphate (0.1%). We determined protein concentration with BCA protein assay kit (ThermoFisher, Waltham, MA, USA) and nomalized Ca content of the cells to protein content.

### 2.10. Quantification of OCN

VICs were cultured in 6-well plates. After removing the medium, we added 100 μL of EDTA (0.5 mol/L, pH 6.9) to the wells. We quantified OCN content of the EDTA-solubilized samples by an enzyme-linked immunosorbent assay (Bio-Techne R&D Systems, Minneapolis, MN, USA). OCN content was normalized to protein content and expressed as ng OCN/mg protein.

### 2.11. Real-time qPCR

RNA was isolated from the hearts of the mice with Tri reagent (Molecular Research Center, Cincinnati, OH, USA) according to the manufacturer’s protocol. To prepare cDNA we used High Capacity cDNA Reverse Transcription Kit (Applied Biosystems, Waltham, USA). The qPCR was carried out on a BioRad CFX96 Real-time System (Bio-Rad, Hercules, CA, USA) with the use of iTaq™ Universal SYBR^®^ Green Supermix (Bio-Rad) and predesigned primers to detect mRNA levels of HIF-1α, HIF-2α, Runx2 and Sox9 (Table 1). We used the comparative Ct method to calculate the expression level of the transcripts, and mouse HPRT was used for normalization as internal control.

**Table 1.**
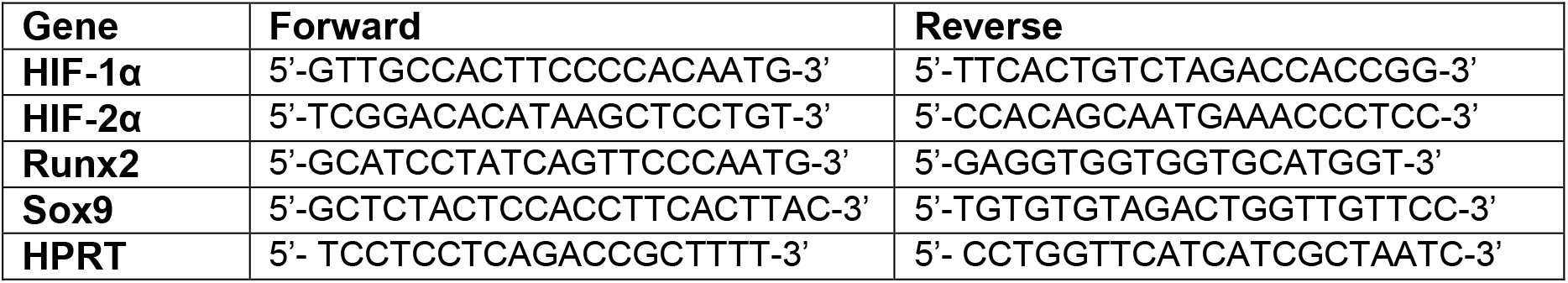
List of primers used in quantitative PCR.

### 2.12. Western blot analysis

We lysed VICs in Laemmli lysis buffer and the cell lysate was resolved by SDS-PAGE (7.5-10%). Proteins were blotted onto nitrocellulose membranes (Amersham, GE Healthcare, Chicago, IL, USA). Western blotting was performed with the use of primary antibodies listed in Table 2. Secondary antibodies – horseradish peroxidase linked rabbit (NA-934) and mouse IgG (NA-931) (Amersham) – were applied at a concentration of 0.5 μg/ml. Blots were developed with enhanced chemiluminescence system Clarity Western ECL (BioRad, Hercules, CA, USA). Chemiluminescent signals were either detected on an X-ray film or with a C-Digit Blot Scanner (LI-COR Biosciences, Lincoln, NE, USA). Following the development, all membranes were stripped and re-probed for β-actin using anti-β-actin antibody at a concentration of 0.5 μg/ml (sc-47778, Santa Cruz Biotechnology Inc., Dallas, TX, USA). We used the inbuilt software of the C-Digit Blot Scanner for quantification.

**Table 2.**
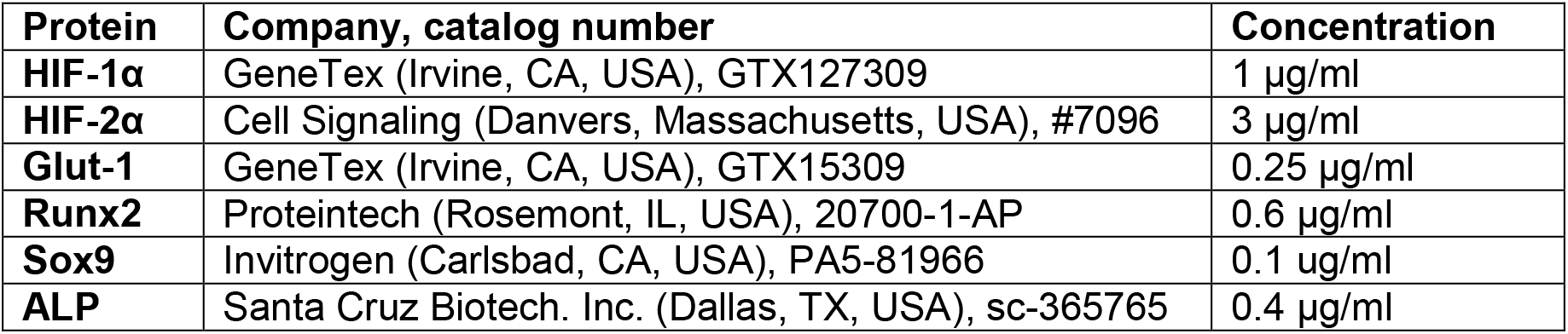
List of primary antibodies used in Western Blot.

### 2.13. RNA silencing

We used Lipofectamine RNAiMAX reagent (Invitrogen, Carlsbad, CA, USA) to transfect VICs with siRNA. We followed the protocol that was provided by the manufacturer. The siRNA for HIF-1α (AM16708, ID: 106498) and HIF-2α (AM16708, ID: 106446) and silencer negative control #1 (4390843) were purchased from Invitrogen. To confirm the efficiency of silencing we performed Western blot analysis.

### 2.14. Intracellular ROS Measurement

The level of ROS was measured with CM-H2DCFDA assay (Life Technologies, Carlsbad, CA, USA). The cells were loaded with the dye (10 μmol/L, 30 minutes), then washed thoroughly with HBSS. After a 4-hour treatment the cells were washed with HBSS and the fluorescence intensity was evaluated with the use of 488 nm excitation and 533 nm emission wavelengths. In some experiments, we applied the ROS inhibitor N-acetyl cysteine (NAC, 1 mmol/L) during the treatment.

### 2.15. Determination of cell viability

We performed MTT assay to measure cell viability. A solution of 3-[4, 5-Dimethylthiazol-2-yl]-2,5-diphenyl-tetrazolium bromide (0.5 mg/mL in HBSS) were incubated with the cells for 4 hours. Following this, we removed the MTT solution and dissolved the formazan crystals in 100 μL of DMSO. Using DMSO as a blank, we measured optical density of the samples at 570 nm.

### 2.16. Data analysis

We show all the results as mean ± SD. We used GraphPad Prism software (version 8.01, San Diego, CA, USA) to perform statistical analyses. Normality of distribution was assessed by Shapiro-Wilk test. All data passed normality and equal variance tests, therefore we used parametric tests to determine p values. Two-tailed Student’s t-test (in case of two groups) and one-way ANOVA followed by Tukey’s post hoc test (in case of more than two groups) were used to determine statistically significant differences between the groups. A value of p < 0.05 was considered significant.

## 3. RESULTS

### 3.1. High phosphate induces heart VC in CKD mice and osteogenic transition of valve interstitial cells (VICs)

Cardiac VC is the main cause of cardiovascular disease and mortality in CKD patients. To set up an *in vivo* model of CKD-associated VC we induced CKD in C57BL/6 mice with a two-phase diet containing adenine (0.2%) and moderately elevated phosphate (0.7%) in the first 6 weeks and adenine (0.2%) and high phosphate (1.8%) in the following 4 weeks. Control mice (Ctrl) received a standard mice diet with 0.3% phosphate content (**Figure 1A**). The development of CKD was accompanied by significant weight loss (**Figure 1B**), and elevation in urea, creatinine and phosphate concentration in plasma (**Figure 1C-E**). To evaluate osteogenic activity in mouse hearts we performed OsteoSense™ staining in Ctrl and CKD mice. Fluorescent intensity of the heart tissue was higher in CKD mice compared to Ctrl mice (4.21×10^8^ vs. 6.99×10^8^ p/s, p<0.001, **Figure 1F**). We performed histological analysis of hearts derived from Ctrl and CKD mice to detect VC. Heart valve of CKD mice showed positive Von Kossa and Alizarin Red staining, whereas no calcification was detectable in the heart of control mice (**Figure 1G**). Additionally, mRNA levels of osteo-, and chondrogenesis-related transcription factors Runx2 and Sox9 were increased in the heart of CKD mice in comparison to Ctrl (**Figure 1H**).

**Figure 1.**
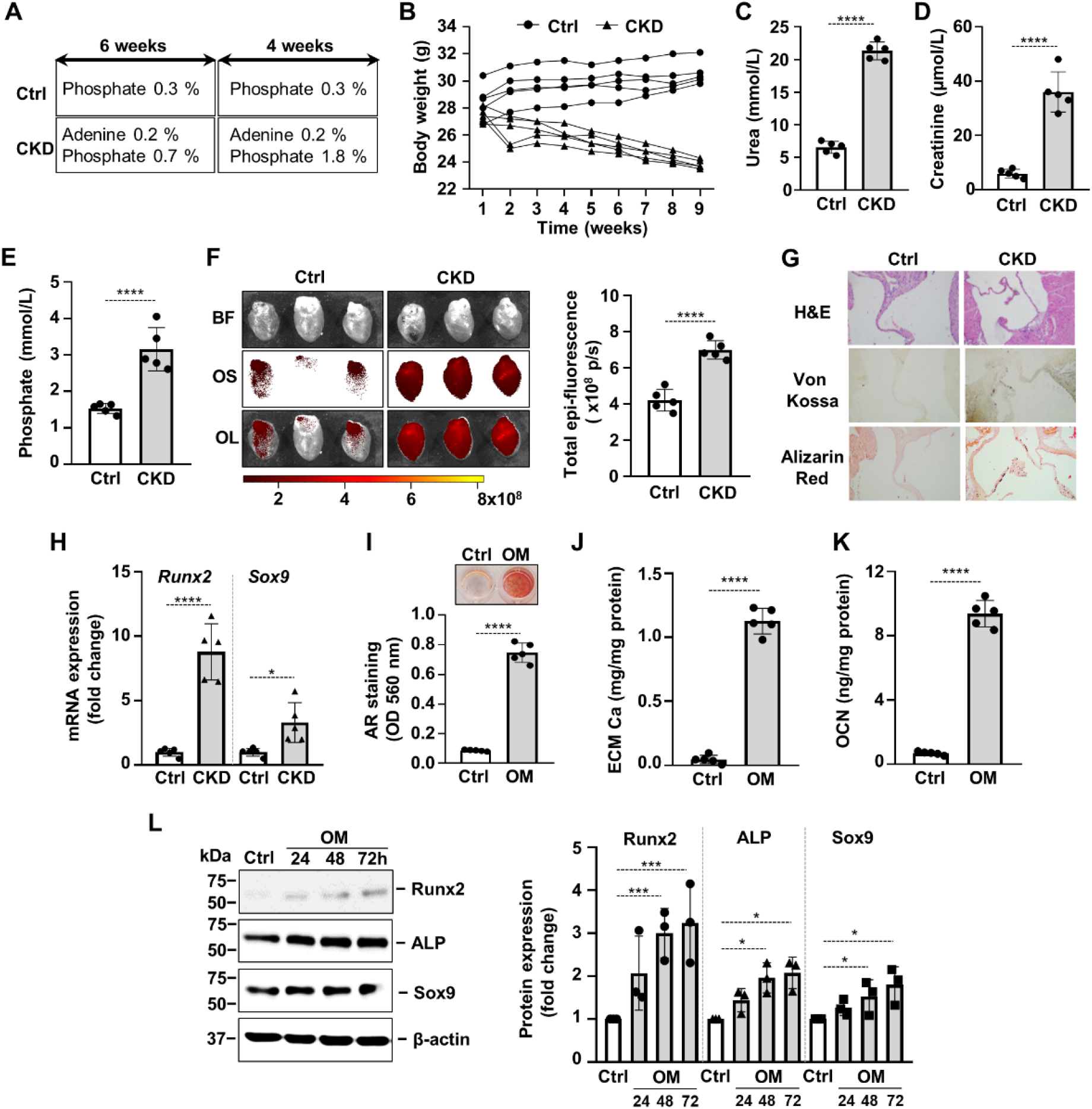
High phosphate induces heart VC in CKD mice and osteogenic transition of valve interstitial cells (VICs). (A) Outline of the experimental protocol. (B) Body weight, (C) plasma urea, (D) plasma creatinine, (E) plasma phosphate levels in control (Ctrl) and CKD mice (n=5/group). (F) Bright-field and macroscopic fluorescence reflectance images of heart and quantification of OsteoSense™ staining intensity in the heart of Ctrl and CKD mice (n=5/group). (G) Representative H&E, von Kossa and Alizarin Red staining of heart sections of Ctrl and CKD mice. (H) Relative mRNA expressions of Runx2 and Sox9 normalized to HPRT from heart tissue derived from Ctrl and CKD mice (n=5, measured in triplicates). Confluent VICs were cultured in Ctrl or osteogenic conditions (OM, 2.5 mmol/L excess Pi, 0.3 mmol/L excess Ca over Ctrl). (I) Calcium deposition in the ECM (day 5) evaluated by AR staining. Representative image and quantification are shown from 5 independent experiments. (J) Calcium content of the HCl-solubilized ECM. (K) OCN level of EDTA-solubilized ECM (day 10). (L) Runx2, ALP and Sox9 protein expressions detected by Western Blot from whole cell lysate (24, 48, 72h). Membranes were re-probed for β-actin. Representative Western blots and densitometry analysis from three independent experiments. Data are expressed as mean ± SD. Ordinary one-way ANOVA followed by Tukey’s multiply comparison test was used to calculate *p* values. \**p*<0.05, \*\*\**p*<0.005, \*\*\*\**p*<0.001

Osteogenic trans-differentiation and extracellular matrix (ECM) mineralization of VICs play a critical role in the progress of cardiac VC. To set up an *in vitro* model of VC we treated VICs with osteogenic medium (OM: growth medium supplemented with 2.5 mmol/L Pi and 0.3 mmol/L Ca). OM triggered calcification of VICs which was assessed by Alizarin Red staining and Ca measurement from HCl-solubilized ECM (**Figure 1I-J**). Furthermore, OM induced deposition of the Ca-binding protein osteocalcin (OCN) in the ECM (**Figure 1K**). In response to OM stimulation we observed time-dependent upregulation of Runx2 and Sox9, as well as alkaline phosphatase (ALP) (**Figure 1L**).

### 3.2. Hypoxia signaling is implicated in high Pi-induced calcification of VICs

Recent works highlighted that hypoxia signaling is activated in calcifying aorta and showed that hypoxia inducible factors (HIFs) play a critical role in osteogenic trans-differentiation of VSMCs ^25–27^. Because vascular and VC share similar mechanisms, next we investigated whether hypoxia signaling is implicated in VC. To this end, first we evaluated the expression of HIF-1α, the regulatory subunit of the HIF-1 transcription factor in heart from Ctrl and CKD mice. As revealed by immunohistochemistry, HIF-1α expression was elevated in the heart valve of CKD mice compared to Ctrl (**Figure 2A**). Additionally, we measured higher mRNA levels of both HIF-1α and HIF-2α in the heart tissue of CKD mice compared to Ctrl (**Figure 2B**). These results suggest that the HIF pathways are activated in the heart of CKD mice.

**Figure 2.**
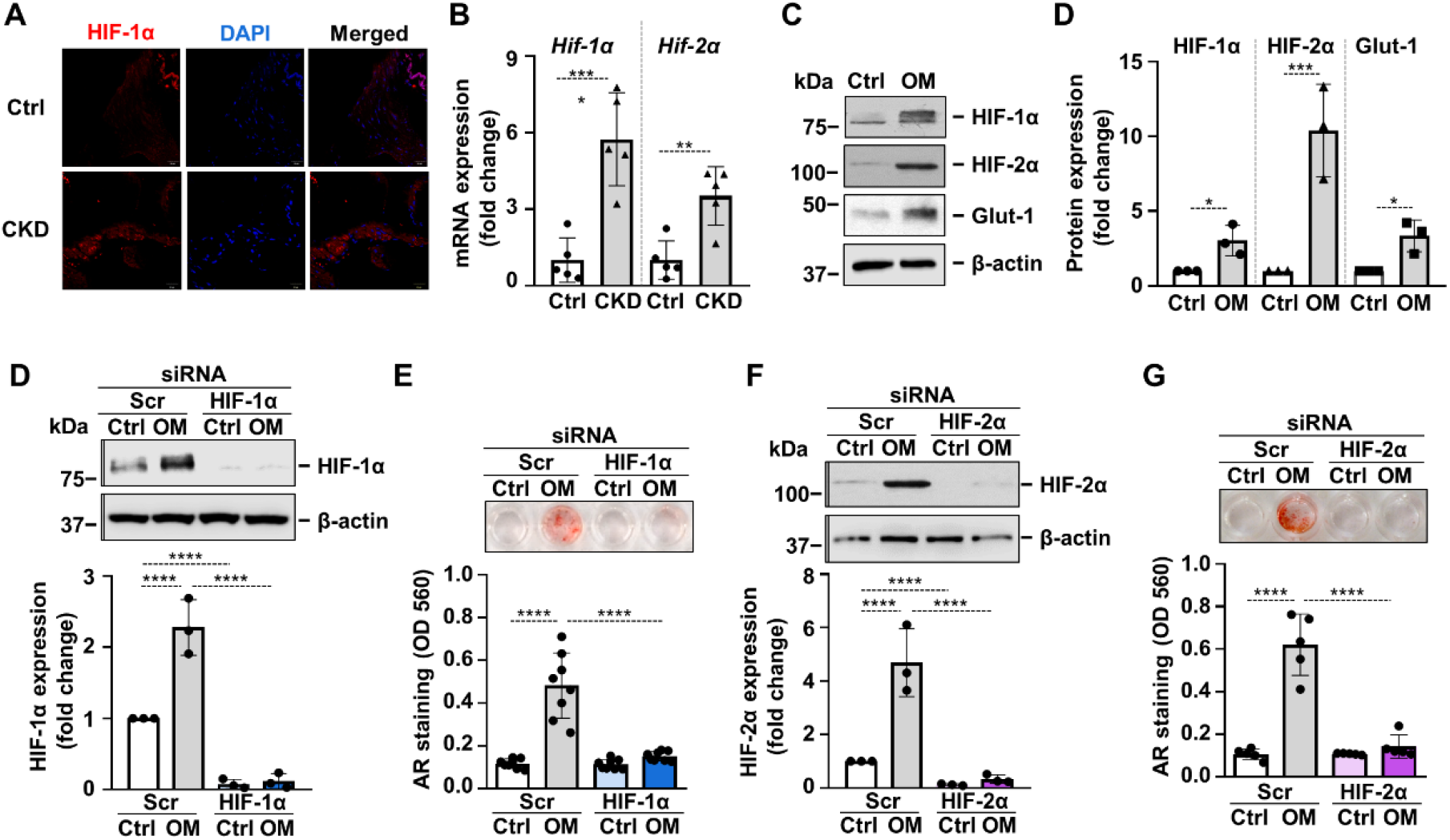
HIF pathway is activated in CKD and is involved in osteogenic trans-differentiation of VICs. (A) Immunohistochemistry was performed on heart tissue samples from Ctrl and CKD mice to detect HIF-1α. Representative HIF-1α (red) and DAPI (blue) staining. (B) Relative mRNA expressions of HIF-1α and HIF-2α normalized to HPRT from heart tissue derived from Ctrl and CKD mice (n=5, measured in triplicates). (C) Confluent VICs were cultured in control (Ctrl) or osteogenic conditions (OM, 2.5 mmol/L excess Pi, 0.3 mmol/L excess Ca over Ctrl). HIF-1α, HIF-2α, Glut-1 and β-actin protein expressions detected by Western Blot from whole cell lysate (24h). Representative Western blots and densitometry analysis from three independent experiments. VICs were kept under Ctrl or OM in the presence of HIF-1α, HIF-2α or scrambled siRNA. (D, F) Protein expression of HIF-1α and HIF-2α in whole cell lysates (24h). Membranes were re-probed for β-actin. Representative Western blots and relative expression of HIF-1α and HIF-2α normalized to β-actin from 3 independent experiments. (E, G) Representative AR staining (day 4) and quantification. Data are expressed as mean ± SD. Ordinary one-way ANOVA followed by Tukey’s multiply comparison test was used to calculate *p* values. \**p*<0.05, \*\**p*<0.001, \*\*\**p*<0.005, \*\*\*\**p*<0.001

Then using our *in vitro* model we showed that osteogenic stimulation with OM triggered a hypoxia response in VICs, characterized by increased protein expressions of HIF-1α, HIF-2α and glucose transporter 1 (Glut-1) (**Figure 2C**). After the establishment that OM triggers HIF activation in VICs, we asked whether this mechanism contributes to the calcification process. To assess this, we applied siRNA to downregulate protein expressions of HIF-1α and HIF-2α, the regulatory subunits of the HIF complexes. Western blots revealed that gene silencing was successful (**Figure 2D and F**). Knockdown of either HIF-1α or HIF-2α was accompanied with decreased calcification of VICs as assessed by Alizarin Red staining (**Figure 2E and G**) suggesting that HIF-1 and HIF-2 pathways are not only activated upon osteogenic stimulation, but they are actively participated in the calcification process.

### 3.3. Hypoxia enhances calcification of VICs in a HIF-1α- and HIF-2α-dependent manner

After defining the crucial involvement of hypoxia signaling in phosphate-induced calcification of VICs we asked whether hypoxia influences OM-induced osteogenic transition and calcification of VICs. First, we exposed VICs to normoxia (21% O_2_) or hypoxia (1% O_2_) and evaluated protein expressions of HIF-1α, HIF-2α and Glut-1. As expected, hypoxia triggered a hypoxia response in VICs characterized by elevated protein expression of HIF-1α, HIF-2α and Glut-1 (**Figure 3A**). Following this, we treated VICs with OM (2.5 mmol/L Pi, 0.3 mmol/L Ca) under normoxic (21% O_2_) and hypoxic (1% O_2_) conditions for 24 and 48 hours. Compared to control, OM treatment slightly increased Runx2 and Sox9 expressions under normoxic condition after 48 hours of exposure (**Figure 3B**). On the other hand, hypoxia strongly upregulated Runx2 expression even in the absence of OM stimulation (**Figure 3B**). Osteogenic stimuli could not further increase Runx2 expression under hypoxia (**Figure 3B**). Compared to normoxia, Sox9 expression was elevated under hypoxia at each condition (**Figure 3B**). These results suggest that hypoxia may exaggerate osteogenic reprogramming of VICs.

**Figure 3.**
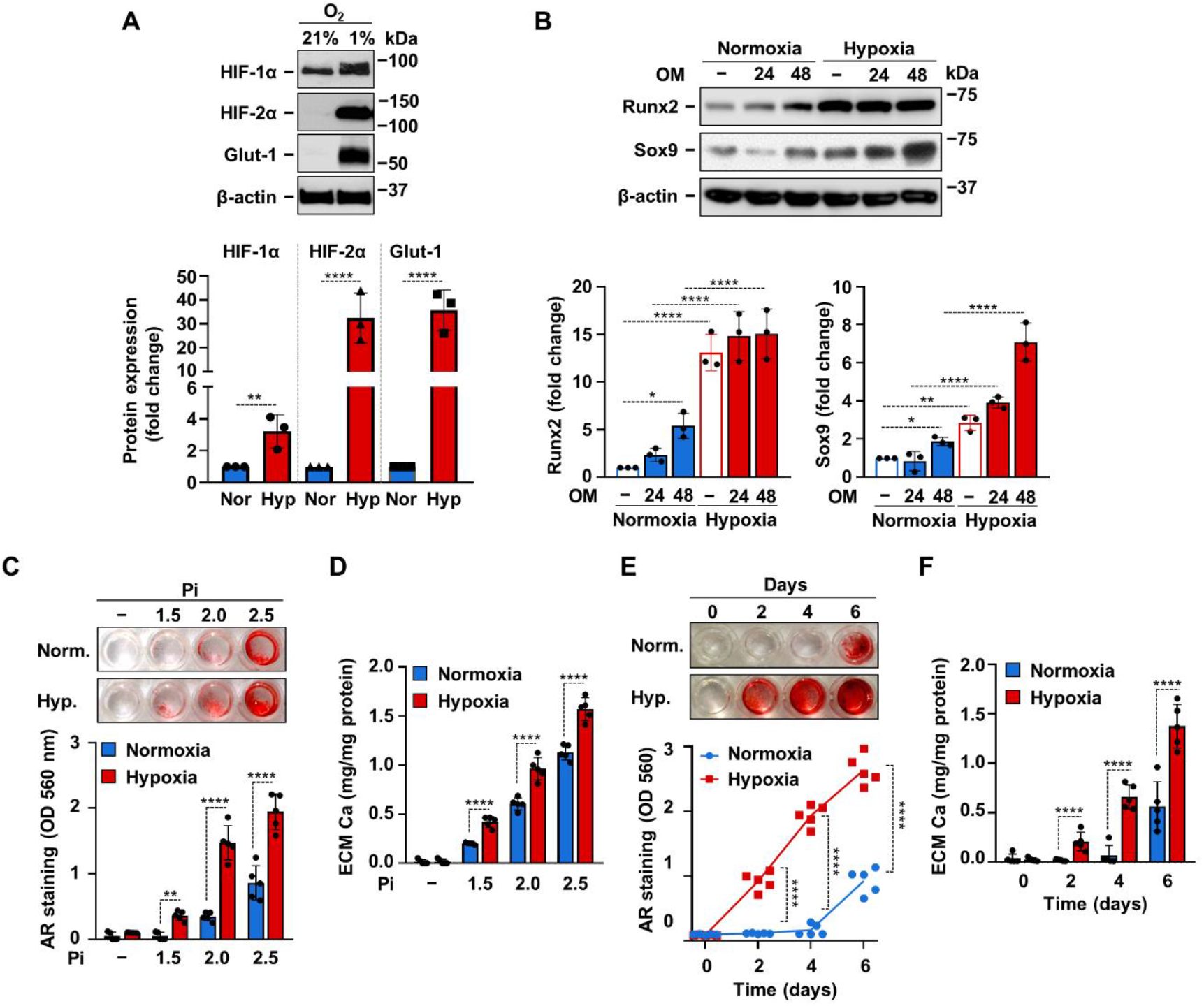
Hypoxia enhances OM-induced calcification of VICs. (A) Confluent VICs were maintained under normoxic (Nor, 21% O_2_) or hypoxic (Hyp, 1% O_2_) conditions. (A) HIF-1α, HIF-2α, Glut-1 and β-actin protein expressions detected by Western Blot from whole cell lysate (24h). Representative Western blots and densitometry analysis from three independent experiments. (B) Confluent VICs under normoxic (21% O_2_) or hypoxic (1% O_2_) conditions were exposed to OM (2.5 mmol/L excess Pi, 0.3 mmol/L excess Ca over Ctrl). Runx2 and Sox9 protein expressions detected by Western Blot from whole cell lysate (24, 48h). Membranes were re-probed for β-actin. Representative Western blots and densitometry analysis from three independent experiments. (C-D) Confluent VICs were exposed to OM with different Pi content (1.5-2.5 mmol/L excess over Ctrl) under normoxic (21% O_2_) and hypoxic conditions (1% O_2_). (C) Representative AR staining (day 6) and quantification. (D) Calcium content of the HCl-solubilized ECM (day 6). (E-F) Time course of calcium accumulation under normoxic and hypoxic conditions in the presence of OM. (E) Representative AR staining and quantification. (F) Calcium content of the HCl-solubilized ECM. Data are expressed as mean ± SD. (A-D and F) Ordinary oneway ANOVA followed by Tukey’s multiply comparison test was used to obtain *p* values. (E) Multiply t-tests to compare normoxia and hypoxia samples at each time points were performed to obtain *p* values. \**p*<0.05, \*\**p*<0.01, \*\*\*\**p*<0.001

Next, we addressed the effect of hypoxia on ECM calcification in VICs. We induced VICs calcification with OM containing calcium (0.3 mmol/L excess) and different amounts of excess Pi (1.5; 2.0; 2.5 mmol/L) under normoxic and hypoxic conditions. As revealed by Alizarin Red staining and calcium measurement hypoxia potentiated the pro-calcification effect of Pi at each tested concentrations (**Figure 3C-D**). Then we investigated time-dependency of VICs calcification under normoxic and hypoxic conditions. Calcification was induced with OM (Ca: 0.3 mmol/L, Pi: 2.5 mmol/L excess), and calcification was investigated on day 2, 4 and 6. Alizarin Red staining showed positivity after 2 days of OM exposure under hypoxic condition, whereas calcification became detectable only on day 6 under normoxia (**Figure 3E**). Calcium measurement from HCl-solubilized ECM also supported the finding that hypoxia potentiates and accelerates Pi-induced calcification of VICs (**Figure 3F**).

To see whether HIF signaling was involved in hypoxia-induced acceleration of VICs calcification, first we applied the HIF inhibitor chetomin and investigated OM-induced calcification under hypoxic condition. As shown by Alizarin Red staining and calcium measurement, chetomin inhibited calcification of VICs (**Figure 4A-B**). Then we knock-down HIF-1α or HIF-2α expressions with the use of target-specific siRNAs under hypoxia. Western blot analysis showed successful knock-down of HIF-1α and HIF-2α protein expressions (**Figure 4C-D**). Silencing of either HIF-1α or HIF-2α resulted in partial inhibition of hypoxia-induced calcification as assessed by Alizarin Red staining (**Figure 4E**) and calcium measurement (**Figure 4F**), supporting the involvement of HIF signaling in hypoxia-induced VICs calcification.

**Figure 4.**
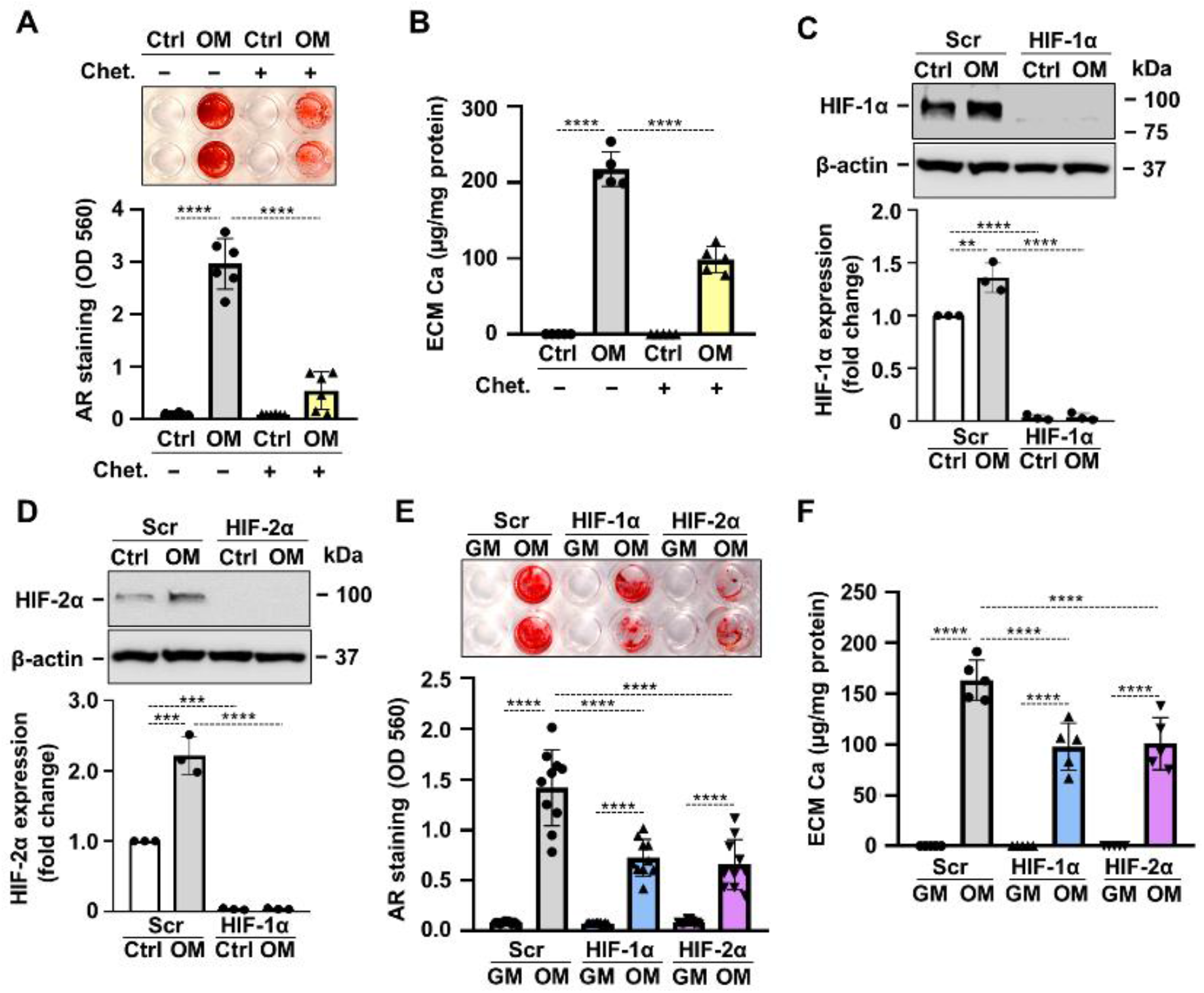
Hypoxia enhances OM-induced osteogenic trans-differentiation of VICs through HIF-1 signaling. (A and B) Confluent VICs were maintained in Ctrl or OM (2.5 mmol/L excess Pi, 0.3 mmol/L excess Ca over Ctrl) conditions under hypoxia (1% O_2_) with or without of the HIF-1 inhibitor chetomin (Chet, 12 nmol/L). (A) Representative AR staining (day 4) and quantification. (B) Calcium content of the HCl-solubilized ECM (day 4). (C-F) VICs were kept under Ctrl or OM conditions in hypoxia (1% O_2_) in the presence of HIF-1α, HIF-2α or scrambled siRNA. (C and D) Protein expressions of HIF-1α and HIF-2α were detected by Western Blot in whole cell lysates (24h). Membranes were re-probed for β-actin. Representative Western blots and densitometry analysis from three independent experiments. (E) Representative AR staining (day 4) and quantification. (F) Calcium content of the HCl-solubilized ECM (day 4). Data are expressed as mean ± SD. *p* values were calculated using one-way ANOVA followed by Tukey’s multiply comparison analysis. \*\**p*<0.001, \*\*\**p*<0.005, \*\*\*\**p*<0.001

### 3.4. The involvement of ROS in hypoxia-mediated potentiation of VICs calcification

Recent evidence suggest a causative role for excess ROS-mediated oxidative stress in the osteogenic differentiation of VICs^13^. There is a multifaceted interplay between hypoxia and ROS formation, eventually leading to excess ROS production under hypoxic conditions. Previously we showed that hypoxia increases VSMCs calcification in a ROS-dependent fashion^25^, therefore our next question was whether unfettered production of ROS is implicated in VICs calcification under hypoxia as well.

To this end, first we measured ROS production in control and OM-stimulated VICs under normoxic and hypoxic conditions. Osteogenic stimulation increased ROS production under normoxia (**Figure 5A**). Compared to normoxia, hypoxia increased ROS production in VICs in both control and OM conditions (**Figure 5A**). The glutathione precursor, N-acetyl-cysteine (NAC) attenuated excessive ROS production in all conditions (**Figure 5A**).

**Figure 5.**
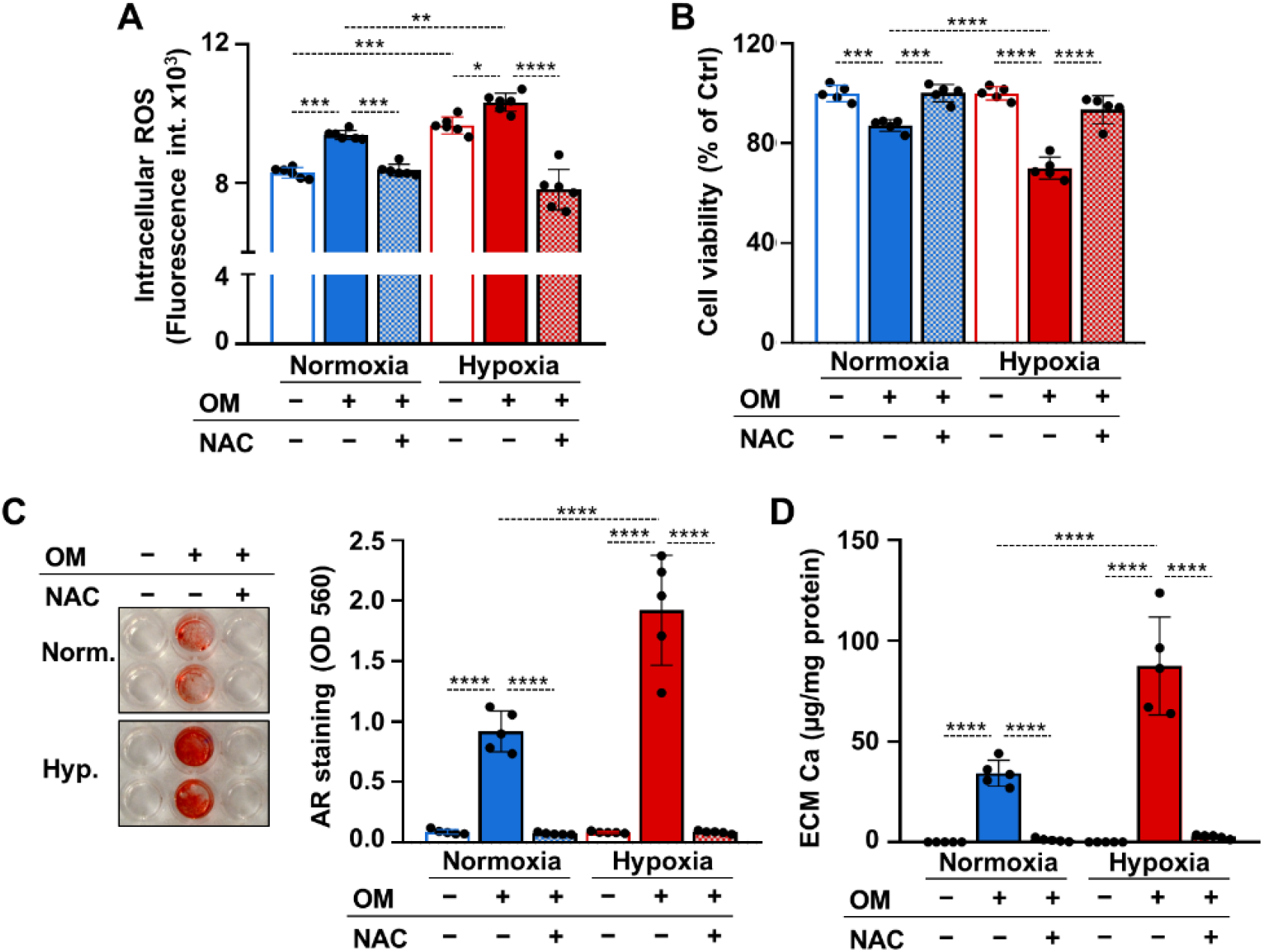
ROS regulate calcification of VICs under both normoxia and hypoxia. (A-D) Confluent VICs were maintained under normoxia (21% O_2_) or hypoxia (1% O_2_) in Ctrl or OM conditions with or without NAC (1 mmol/L). (A) Intracellular ROS formation in VICs (4h). (B) Cell viability assessed by MTT assay (day 4). (C) Representative AR staining (day 4) and quantification. (B) Calcium content of the HCl-solubilized ECM (day 4). Data are expressed as mean ± SD. Ordinary one-way ANOVA followed by Tukey’s multiply comparison test was used to obtain *p* values. \**p*<0.05, \*\**p*<0.01, \*\*\**p*<0.005, \*\*\*\**p*<0.001

Apoptotic cell death and the release of apoptotic bodies is an important calcification mechanism. Excess ROS production can trigger cell death, therefore next we investigated cell viability in control and OM-treated VICs under normoxia and hypoxia after 4 days of exposure in the presence or absence of NAC. Osteogenic stimulation triggered a decline in cell viability in normoxia and even more cell death was observed in hypoxia (**Figure 5B**). NAC prevented OM-induced cell death under both normoxia and hypoxia (**Figure 5B**). Attenuation of unfettered ROS production and cell death by NAC was associated with complete inhibition of OM-induced VICs calcification as revealed by Alizarin red staining and calcium measurements (**Figure 5C and D**).

### 3.5. Chemical hypoxia mimetic drugs enhance VICs calcification

Hypoxia mimetic drugs mimic the effect of real hypoxia through the stabilization of HIFα subunits. We investigated three different hypoxia mimetic drugs, cobalt-chloride (CoCl2), desferrioxamine (DFO) and Daprodustat (DPD), to see whether they influence Pi-induced VICs calcification under normoxic condition. We treated VICs with CoCl2 (200 μmol/L), DFO (40 μmol/L) or DPD (20 μmol/L) for 24 hours and first we evaluated protein expressions of HIF-1α and HIF-2α from whole cell lysate (**Figure 6A**). As expected, hypoxia mimetics stabilized both HIF-1α and HIF-2α subunits in VICs.

**Figure 6.**
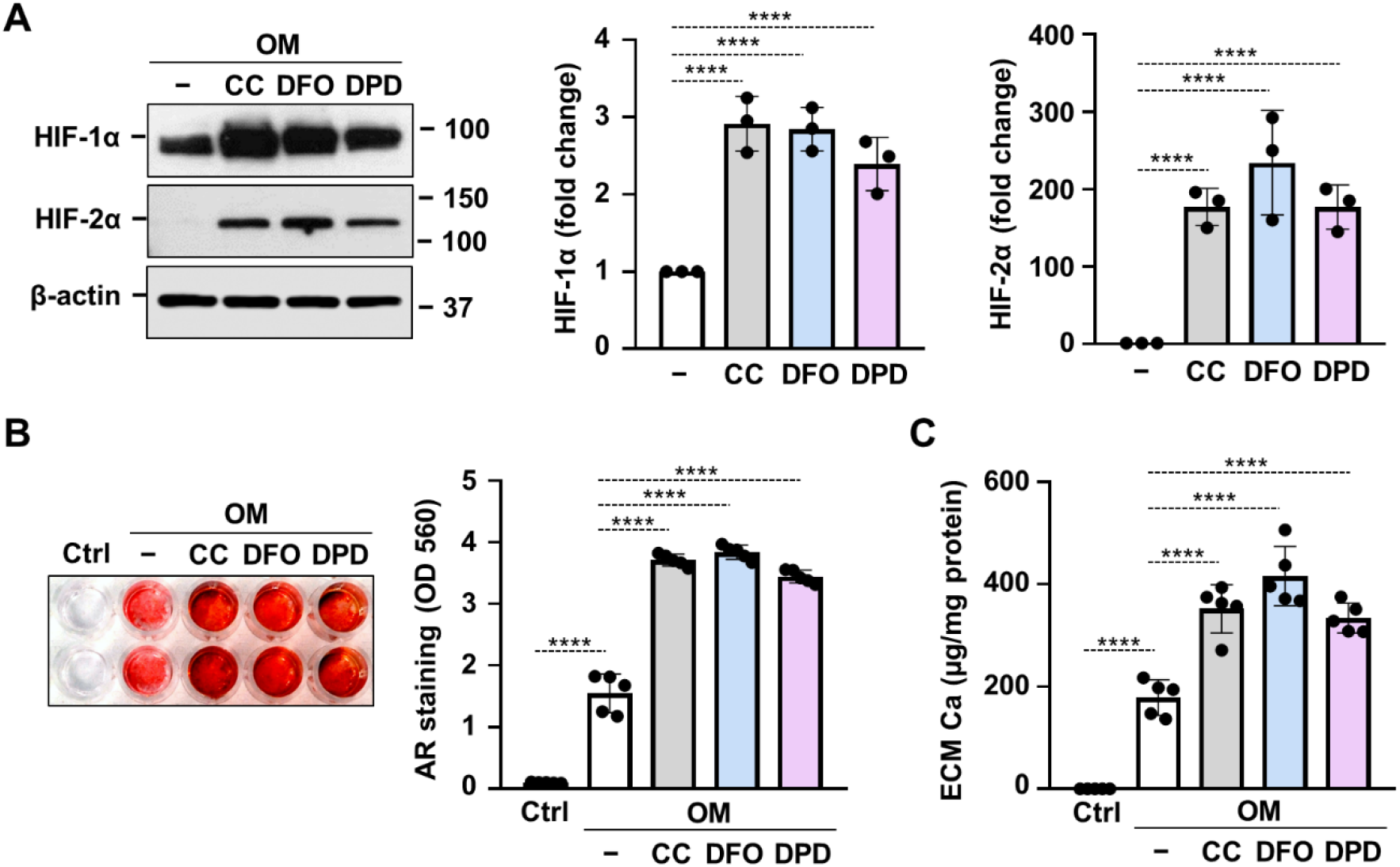
Hypoxia mimetic drugs augment OM-induced calcification of VICs. (A-C) Confluent VICs maintained in OM (2.5 mmol/L excess Pi, 0.3 mmol/L excess Ca) were treated with hypoxia mimetic drugs CoCl2 (CC, 200 μmol/L), desferrioxamine (DFO, 40 μmol/L) and Daprodustat (DPD, 20 μmol/L). (A) Protein expressions of HIF-1α and HIF-2α were detected by Western Blot in whole cell lysates (24h). Membranes were re-probed for β-actin. Representative Western blots and densitometry analysis from three independent experiments. (B) Representative AR staining (day 5) and quantification. (C) Calcium content of the HCl-solubilized ECM (day 5). Data are expressed as mean ± SD. Ordinary one-way ANOVA followed by Tukey’s multiply comparison test was used to obtain *p* values. \**p*<0.05, \*\**p*<0.01, \*\*\**p*<0.005, \*\*\*\**p*<0.001

Next, we addressed whether hypoxia mimetic drugs influence OM-induced calcification of VICs. We treated VICs with OM (0.3 mmol/L excess Ca, 2.5 mmol/L excess Pi) in the presence or absence of CoCl2 (200 μmol/L), DFO (40 μmol/L) or DPD (20 μmol/L). Alizarin Red staining and calcium measurement were performed on day 5. We observed that all the three tested hypoxia mimetic drugs enhanced OM-induced calcification in VICs (**Figure 6B and C**). These results suggest that not only real hypoxia but also chemical activation of the HIF pathways enhances calcification of VICs.

### 3.6. DPD enhances aortic VC in CKD mice

DPD is a hypoxia mimetic drug that is used to treat anemia in CKD patients in Japan. After seeing that DPD enhances VICs calcification *in vitro* we addressed its effect on VC in the adenine-induced CKD model. Fifteen C57BL/6 mice (8-10 weeks old, male) were assigned to 3 groups, Ctrl, CKD, and CKD+DPD (**Figure 7A**). DPD was administered at the dose of 15 mg/body weight kg/day orally between week 7 and 10 (**Figure 7A**). Anemia was developed in CKD mice, characterized by decreased Hb concentration, and low red blood cell count and hematocrit levels (**Table 3**). On the contrary, Hb concentration, red blood cell count and hematocrit levels were normal in CKD mice with DPD treatment (**Table 3**).

**Figure 7.**
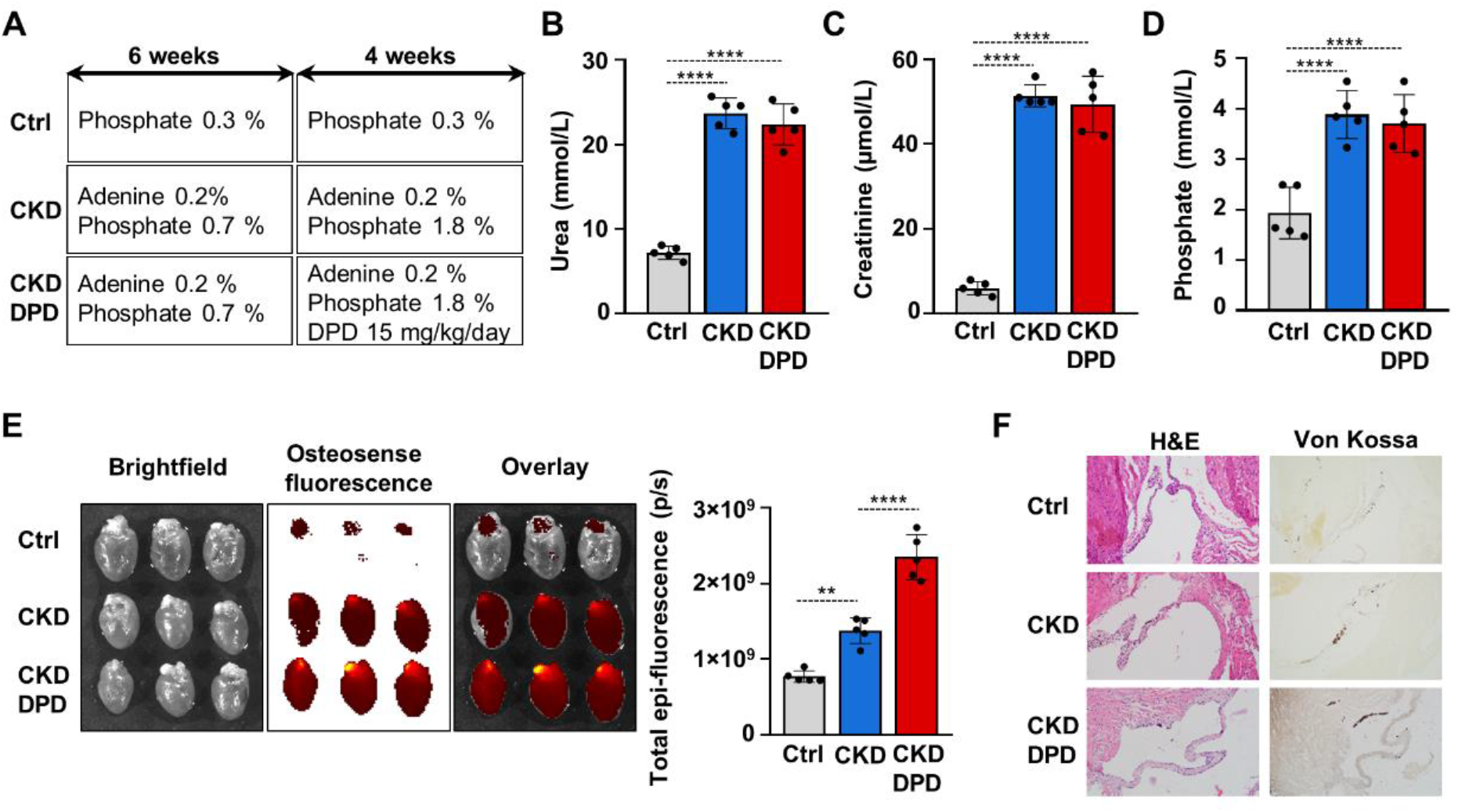
DPD increases aortic VC in mice with CKD. (A) Outline of the experimental protocol. (B) Plasma urea, (C) creatinine, (D) phosphate levels (n=5/group). (E) Bright-field and macroscopic fluorescence reflectance images of heart and quantification of OsteoSense™ staining intensity in the heart of Ctrl, CKD and CKD+DPD mice (n=5/group). (F) Histological analysis of heart valves derived from Ctrl, CKD, and CKD+DPD mice. Representative H&E and von Kossa-stained heart sections. Data are expressed as mean ± SD. Ordinary one-way ANOVA followed by Tukey’s multiply comparison test was used to obtain *p* values. \*\**p*<0.01, \*\*\*\**p*<0.001

**Table 3.** Hematology parameters.

| Hematology parameter | Control | CKD | CKD+DPD | *p* value Ctrl vs CKD | *p* value CKD vs CKD+DPD |
| --- | --- | --- | --- | --- | --- |
| Hemoglobin (g/L) | 117.4 ± 3.6 | 80.2 ± 8.9 | 119.2 ± 3.3 | 0.000012 | 0.0000077 |
| Red blood cell count (T/L) | 8.014 ± 0.15 | 6.044 ± 0.561 | 8.046 ± 0.29 | 0.000032 | 0.000052 |
| Hematocrit | 0.439 ± 0.017 | 0.301 ± 0.031 | 0.432 ± 0.014 | 0.000012 | 0.000013 |

On the other hand, plasma urea, creatinine and phosphate levels were similarly high in DPD- and vehicle-treated CKD mice (**Figure 7B-D**). To address the effect of DPD on heart calcification we performed OsteoSense™ staining and detected higher amount of hydroxyapatite deposition in the hearts derived from DPD-treated CKD mice compared to vehicle-treated CKD mice (2.35×10^9^ ± 0.3×10^9^ vs. 1.38×10^9^ ± 0.17×10^9^ p/s, p<0.05) (**Figure 7E**). Additionally, we performed histological analysis of hearts derived from Ctrl, CKD and CKD+DPD mice to detect VC. We found stronger von Kossa staining in heart valves of CKD+DPD mice compared to CKD, whereas no calcification was detectable in the heart of Ctrl mice (**Figure 7F**). These results suggest that DPD - at the dose that is efficient to correct CKD-associated anemia -, can accelerate VC in mice with CKD.

## 4. DISCUSSION

A declining kidney function is associated with a significantly reduced life expectancy for both genders in all age groups ^29,30^. The higher mortality associated with renal impairment is mainly due to the increased risk of cardiovascular disease (heart failure and valvular disease)^31^. VC is a frequent and early complication in patients on hemodialysis which contributes to the occurrence of major adverse cardiac events^1–3^.

Deterioration in renal phosphate clearance begins at early stages of CKD which manifests as hyperphosphatemia as the disease progresses^32^. Higher serum phosphate concentrations were found to be linked with an increased prevalence of VC in patients with moderate CKD^33^. In response to high phosphate, VICs, the major cell type composing valves, undergo an osteogenic trans-differentiation process which is thought to be involved in VC.

Here we first established *in vivo* and *in vitro* models of high Pi-induced VC and osteogenic trans-differentiation of human VICs. We used the adenine-induced CKD mice model which was developed originally by Tani et al. to study the progression of hyperphosphatemia and associated mineral bone disease^28^. Adenine+high Pi diet induced CKD in C57BL/6 mice characterized by elevated plasma urea and creatinine levels (**Figure 1C and D**), and hyperphosphatemia (**Figure 1E**). Additionally, we presented that adenine+high Pi diet induced hydroxyapatite accumulation in the heart valves (**Figure 1F-G**), showing that this model is suitable to study Pi-induced VC. In agreement with our results using the same mice model Dargam et al. recently detected VC by S2 heart sound^34^. The synthetic bisphosphonate fluorescence dye OsteoSense™ that binds specifically to hydroxyapatite, is getting more recognition in the detection of vascular calcification^27,35^. Here we succeeded to use OsteoSense™ to assess hydroxyapatite deposition in the heart tissue (**Figure 1F**).

Genome-wide association studies highlighted the role of Runx2, the key transcription factor associated with osteoblast differentiation, as a possible driver of calcific aortic valve stenosis^36^. The causal role of Runx2 in the development of aortic stenosis was also supported by the finding that conditional depletion of Runx2 in mouse models of calcific aortic valve disease resulted in decreased osteogenic gene expression and attenuation of aortic VC^37^. Here we showed marked upregulation of Runx2 mRNA in the heart of CKD mice compared to controls (**Figure 1H**), and elevation of Runx2 protein expression in VICs exposed to high Pi-containing OM (**Figure 1L**). Sox9, the master transcription factor of chondrogenesis, is largely implicated in the mechanism of vascular calcification through promoting chondrogenic differentiation of VSMCs^38^. On the other hand, Sox9 in heart valves prevents matrix mineralization, and reduced Sox9 function is associated with heart VC^39,40^. Our results revealed high basal expression of Sox9 in VICs which was slightly upregulated by osteogenic stimulation (**Figure 1L**).

Tissue hypoxia is concerned in the pathomechanism of many human diseases including kidney disease ^41,42^. Hypoxia accelerates the progression of CKD via promoting fibrogenesis of renal fibroblasts, and triggering epithelial-mesenchymal transformation of renal tubular cells^43,44^. Due to CKD-associated anemia and damage of the microvasculature, tissue hypoxia in CKD is not limited to kidney but affects other organs as well^45,46^. In line of this notion, here we showed increased mRNA and protein expression of HIF-1α and HIF-2α in heart derived from CKD mice (**Figure 2A-B**).

Mokas et al. previously reported that high Pi triggers a hypoxic response in VSMCs and proved that HIF-1α-mediated signaling plays a critical role in Pi-induced VSMCs calcification^26^. Here we found that a similar mechanism is present in VICs; Pi triggers activation of the HIF pathway which is necessary for the Pi-induced calcification of VICs (**Figure 2C-G**). Pi is a strong inducer of Runx2 in diverse cells and previous studies indicated that Runx2 competes with von Hippel Lindau protein for binding to HIF-1α and blocks the ubiquitination and subsequent degradation of HIF-1α^47^. The potential relevance of this mechanism in Pi-induced stabilization of HIF-1α and HIF-2α in VICs needs to be investigated.

Further activation of the HIF pathways by hypoxia largely promotes Pi-induced calcification of VICs (**Figure 3**). Similar stimulating effect of hypoxia on osteogenic differentiation of VSMCs, multipotent human mesenchymal stromal cells and periosteal cells have been reported^26,48,49^. In a recent study Kanno et al. showed elevated expression of numerous mesenchymal and hematopoietic progenitor markers in VICs under hypoxic (2% O_2_) culture conditions, and connected stemness of hypoxic VICs with increased potential towards osteogenic differentiation^50^.

Opposing to our results, hypoxia (2% O_2_) has been shown to decrease the expression of osteogenic markers in MG63 osteoblast-like cells^51^. According to another study, hypoxia (2% O_2_) does not influence osteogenic differentiation of primary osteoblasts and mesenchymal precursors, but quick exposure to anoxia inhibits bone nodule formation and calcification through the downregulation of Runx2^52^. Overall, these results suggest that the effect of hypoxia on osteogenic differentiation is finely regulated and cell specific, in which the differences in Runx2 promoter activity in osseous and non-osseous cells might play a role^53^.

Cellular hypoxic response is mediated by the heterodimeric transcription factors HIF-1 and HIF-2 which are regulated by their oxygen-sensitive alpha subunits HIF-1α and HIF-2α, respectively. HIF-1α and HIF-2α shows 48% amino acid sequence homology and the proteins have similar domain arrangements^54^. Besides that, both HIF-1α and HIF-2α dimerize with HIF-1β and the heterodimer binds to hypoxia response elements of hypoxia-regulated genes^54^. On the other hand, while HIF-1α is expressed ubiquitously, the expression of HIF-2α varies in different tissues, which gives a rise to cell-specific functions of HIF-1 and HIF-2^54,55^. In the heart both HIF-1 and HIF-2 pathways are active, but their particular functions are not fully understood. During hypoxia HIF-1 regulates glucose metabolism and play a crucial role in regulating the angiogenic response following myocardial injury, while the role of HIF-2 remained to be defined^56^. Here we showed that both HIF-1 and HIF-2 pathways are involved in osteogenic differentiation of VICs under both normoxic and hypoxic conditions (**Figure 2, Figure 4**). A direct physical interaction between HIF-1α and Runx2, and its function in regulating angiogenic response in pre-osteoblast cells have been described^57^. Additionally, HIF-2α was identified as a strong activator of Runx2 promoter activity^53^ but the involvement of these mechanism in hypoxia-induced promotion of VICs calcification remained to be elucidated.

At physiological concentrations, ROS regulate major cellular functions such as cell growth and differentiation. On the other hand, excess ROS formation has been implicated in various disorders including cardiovascular disease. Growing evidence highlighted the important causative role for ROS in vascular calcification and calcific aortic valve disease ^13,58^. In line of this notion, increased ROS production was detected in aortic valve tissue from patients with pathological heart valve dysfunctions when compared with transplant-derived control tissues^59^. Additionally, elevated ROS level was associated with decreased expression of antioxidant enzymes such as catalase and all isoforms of superoxide dismutase (SOD1-3)^59^. In the same study authors showed that hydrogen-peroxide induced osteogenic differentiation of VICs which was characterized by increased expression of Runx2 and decreased expression of alpha smooth muscle actin^59^.

The relation between hypoxia and ROS production is controversial, but majority of the evidence suggests that hypoxia stimulates ROS formation in most types of mammalian cells^60^. Hypoxia impairs the function of the mitochondrial electron transport chain complexes leading to increased ROS signals that trigger hypoxia response in diverse cell types^61–63^. Additionally, a study on pulmonary artery smooth muscle cells revealed that hypoxia-induced mitochondrial ROS activates NADPH oxidases which provides a positive feedback loop of exacerbated ROS formation upon hypoxia^64^.

Here we found that hypoxia increases ROS formation in both control and OM-treated cells (**Figure 5A**). OM stimulation under hypoxia reduced cell viability substantially (**Figure 5B**). Inhibition of ROS production by NAC protected VICs from OM-induced cell death under hypoxia, supporting the causative role of ROS in this process (**Figure 5A and B**). The role of programmed VSMCs death is well-established in the pathomechanism of atherosclerosis and vascular calcification ^65,66^. According to our results NAC also inhibited OM-induced calcification of VICs under both normoxic and hypoxic conditions, suggesting the involvement of exacerbated ROS production in the calcification process (**Figure 5C and D**).

Activation of the HIF pathways takes place through stabilization of the HIF α subunits. Normally, HIF α subunits are hydroxylated at specific proline residues by prolyl hydroxylase domain proteins (PHDs) and eliminated via the ubiquitin-proteasome degradation pathway^67^. PHD-mediated prolyl hydroxylation requires oxygen, and dependent on iron, 2-oxoglutarate, and ascorbic acid. Depletion of any of these factors compromise PHD activity. Here we tested three different PHD inhibitors, cobalt chloride that depletes ascorbic acid, DFO, an iron chelator and Daprodustat (DPD), a small molecule inhibitor developed by GlaxoSmithKline^67–69^. All the three PHD inhibitors stabilized both HIF-1α and HIF-2α subunits and promoted OM-induced calcification of VICs under normoxic condition (**Figure 6**). In agreement with this result previous studies showed enhancement of Pi-induced calcification by DPD and Roxadustat in VSMCs^27,70^.

CKD is often accompanied by other chronic conditions such as anemia^71^. Anemia of CKD patients was treated with recombinant erythropoietin or erythropoiesis-stimulating agents (ESAs)^72^. Unfortunately, recent studies showed that ESAs increase the risks for major cardiovascular events and accelerate disease progression in CKD patients ^73–76^. This urged the development of alternative therapeutic approaches. DPD is a new-generation drug and approved in Japan since 2020 for the treatment of patients with CKD-associated anemia ^77,78^.

After seeing that DPD promoted calcification of VICs *in vitro,* we investigated the effect of DPD on VC in a mice model of CKD. Our results revealed that DPD corrects anemia as was described previously, but unfortunately, the beneficial effect of DPD comes at a price as DPD promoted VC *in vivo* (**Figure 7**). Previously we found similar effect of DPD on aorta calcification^27^. Recent phase 3 trials compared the effect of DPD and an injectable ESA in anemic (Hb: 8.0-11.5 g/dL) dialyzed and non-dialyzed patients with CKD^79,80^. These two trials concluded that DPD was non-inferior to ESA with respect to the increase in the Hb level from baseline in both dialysis-dependent and dialysis-independent CKD patients^79,80^. Additionally, they found that the percentages of patients with adverse cardiovascular events were similar in the DPD and ESA groups among CKD patients regardless of dialysis status^79,80^.

In conclusion, here we showed that hypoxic or pharmacological activation of the HIF pathway accelerates phosphate-induced calcification of VICs, in a HIF-1α, HIF-2α and ROS-dependent manner. The new generation PHD inhibitor DPD increased VC *in vivo* in a murine CKD model with high plasma phosphate level. Further studies are needed to investigate whether this mechanism contributes to the occurrence of major cardiovascular events which was reported to happen in 25.2% of hemodialysis-dependent CKD patients on DPD treatment during a 2.5-year follow-up period, and in 19.5% of non-dialyzed CKD patients on DPD treatment during a 1.9-year follow-up period^79,80^.

## Funding statement

This work was funded by the Hungarian National Research, Development and In-novation Office (NKFIH) [K131535 to V.J., FK135327 to B.N. and K139396 to T.J.]; the Hungarian Academy of Sciences [MTA-DE Lendület Vascular Pathophysiology Research Group, grant number 96050 to V.J.]. E.B. was supported by the János Bolyai Research Scholarship of the Hungarian Academy of Sciences. E.B. and A.T. were supported by the New Excellence Program of the Ministry of Human Capacities of Hungary.

## Author contribution statement

VJ designed the research; DMC, HA, AT, GL, ÁS, AT, CF, TJ, BN, EB and VJ performed the experiments; VJ, DMC, AT, ÁS, TJ, BN, and EB analyzed and interpreted the data; and VJ wrote the manuscript. The manuscript was reviewed and edited by every author.

## Conflict of interest statement

The authors declare no conflicts of interest.

## Abbreviations

ALP: alkaline phosphatase
AR: alizarin red
BMP2: bone morphogenetic protein 2
CKD: chronic kidney disease
Ctrl: control
DFO: Desferrioxamine
DMEM: Dublecco’s modified eagle medium
DMSO: dimethyl sulphoxide
DPBS: Dulbecco’s phosphate-buffered saline
DPD: Daprodustat
ECM: extracellular matrix
EPO: erythropoietin
ESAs: erythropoiesis-stimulating agents
FBS: fetal bovine serum
Glut-1: glucose transporter 1
HIF: hypoxia inducible factor
H&E: hematoxylin eosin
NAC: N-acetyl cysteine
OCN: osteocalcin
OD: optical density
OM: osteogenic medium
OPN: osteopontin
Pi: inorganic phosphate
ROS: reactive oxygen species
Runx2: Runt-related transcription factor 2
VC: valve calcification
VICs: valve interstitial cells
VSMCs: vascular smooth muscle cells

## Notes

### Competing Interest Statement

The authors have declared no competing interest.

